# Nucleosome-bound NR5A2 structure reveals pioneer factor mechanism by minor groove anchor competition

**DOI:** 10.1101/2023.06.14.544949

**Authors:** Wataru Kobayashi, Anna Sappler, Daniel Bollschweiler, Maximilian Kümmecke, Jérôme Basquin, Eda Nur Arslantas, Siwat Ruangroengkulrith, Renate Hornberger, Karl Duderstadt, Kikuë Tachibana

## Abstract

Gene expression during natural and induced reprogramming is controlled by pioneer transcription factors, which bind nucleosomal DNA and initiate gene expression from closed chromatin. How pioneer factors perform this function is poorly understood. Nr5a2 is a pioneer factor that is critical for zygotic genome activation (ZGA) in totipotent embryos, pluripotency in embryonic stem cells and metabolic regulation in adult tissues. To study how Nr5a2 functions as a pioneer factor, we used cryo-electron microscopy to determine a structure of human NR5A2 bound to a nucleosome. The structure shows that the conserved C-terminal extension (CTE) loop of the NR5A2 DNA-binding domain competes with a DNA minor groove anchor of the nucleosome and releases entry-exit site DNA at a ∼50° angle. Residue aspartic acid 159 of the CTE loop induces the rearrangement by shifting a histone H2A α2 helix towards the nucleosome center. Mutational analysis showed that NR5A2 D159 is largely dispensable for DNA binding but required for stable nucleosome association and persistent DNA unwrapping. These findings suggest that NR5A2 belongs to a previously unknown class of pioneer factors that can use minor groove anchor competition to destabilize nucleosomes and facilitate gene expression changes during reprogramming.

## Introduction

Gene regulation by transcription factors is important for cell identity. Transcription factors scan chromatin and bind to their cognate motifs in *cis*-regulatory elements such as promoters and enhancers ^1^. DNA access is restricted by nucleosomes that can be inhibitory for transcription factor binding ^2,3^. Transcription factors with pioneer activity, hereafter referred to as pioneer factors, can bind to their (partial) motif on the nucleosome and elicit local opening of closed chromatin ^4-6^. The first transcription factor to be designated as a pioneer factor, FoxA1, functions by replacing the linker histone H1 ^7,8^. Recent structural studies of nucleosomes bound by pluripotency-related pioneer factors of the OCT4 and SOX family members showed several binding modes and structural changes in the nucleosome that depended on the motif position on the nucleosome ^9-11^. Binding of other pioneer factors like GATA3 does not alter nucleosome architecture ^12^. Different pioneer factors might therefore use different mechanisms to remove DNA from nucleosomes but the molecular basis of this function is not well understood.

Whilst several pioneer factors interact with the major groove of DNA, certain nuclear receptors show dual sequence recognition by binding major and minor grooves at the same time ^13^. A notable pioneer factor belonging to this class is the orphan nuclear receptor Nr5a2 (also known as Liver hormone receptor 1, Lrh-1), which has essential functions in development, health and disease ^14-17^. Nr5a2 regulates gene expression in adult tissues including liver and pancreas, and its haploinsufficiency is linked to gastrointestinal and pancreatic cancer ^18^. Nr5a2 is essential for maintaining naïve pluripotency in mouse embryonic stem cells ^19^. Recently, Nr5a2 has been identified as a key regulator of zygotic genome activation in murine embryos ^20^. Interestingly, Nr5a2 can replace Oct4 to potently improve the efficiency of induced reprogramming of somatic cells to pluripotency ^21^.

How NR5A2 engages with nucleosomes to facilitate extreme gene expression changes such as during natural and induced reprogramming is not known. The protein consists of a putative ligand binding domain (LBD), whose ligand is unknown, and a DNA binding domain (DBD) that is conserved throughout metazoa ^22^ (Extended Data Fig.1a). The DBD consists of two modules of Cys_4_-type zinc fingers, a C-terminal extension (CTE), and a Ftz-f1 (fushitarazu-factor 1) domain (Extended Data Fig.1b,c). A crystal structure of the NR5A2-DNA complex showed that the α-helix of the zinc finger domain and the CTE bind to the major and minor groove of DNA, respectively ^23^(Extended Data Fig.1b). Unlike many other nuclear receptors, Nr5a2 binds as a monomer to its DNA motif TCAAGG(C/T)CA” on naked DNA and nucleosomes ^20,23,24^. But how orphan nuclear receptors such as Nr5a2 engage with nucleosomes for pioneer activity is not known.

## Results

### Structure of NR5A2-nucleosome complex shows unwrapping of DNA from histones

To investigate the mechanism of NR5A2 pioneer function, we performed Selected Engagement on Nucleosome Sequencing (SeEN-seq) to identify the preferred binding site on the nucleosome to generate a homogeneous population of protein complexes for cryo-EM ^10^. The Nr5a2 motif was tiled at 1 bp intervals through a 601-nucleosome positioning sequence to simultaneously interrogate all potential transcription factor binding registers ^25^ (Extended Data Fig.2a-d). Electrophoretic mobility shift analysis (EMSA) showed band shifts corresponding to human NR5A2 or mouse Nr5a2 with nucleosome libraries (Extended Data Fig.2e,f). Both preferentially targeted the entry-exit sites on the nucleosome, as was shown previously using 5 bp intervals ^20^ (Fig. 1a, Extended Data Fig.2g,h, and Table 1). Both orthologues displayed a similar binding pattern with a ∼10 bp periodicity (e.g. motif positions 10, 20, 30, 40) that reflects (partial) motif accessibility on the rotational turns of the DNA double helix (Fig 1a, and Extended Data Fig.2g,h). Since NR5A2 showed a stronger binding site preference than Nr5a2, we proceeded with the human protein.

**Table 1:** Cryo-EM data collection, refinement and validation statistics.

|  | NR5A2-nucleosome complex<sup>SHL+5.5</sup><br>(EMDB-xxxx)<br>(PDB xxxx) | 601nucleosome<sup>SHL+5.5</sup><br>(EMDB-xxxx)<br>(PDB xxxx) | 601nucleosome<sup>SHL+5.5</sup><br>(EMDB-xxxx) |
| --- | --- | --- | --- |
| **Data collection and processing** |  |  |  |
| Magnification | 105000 | 105000 | 22000 |
| Voltage (kV) | 300 kV | 300kV | 200 kV |
| Electron exposure (e-/Å<sup>2</sup>) | 65.4 | 65.4 | 65 |
| Defocus range (µm) | -0.6 - -2.2 | -0.6 - -2.2 | -1 - -2.6 |
| Pixel size (Å) | 0.8512 | 0.8512 | 1.885 |
| Symmetry imposed | none | none | C2 |
| Initial particle images (no.) | 3,109,326 | 3,109,326 | 463,334 |
| Final particle images (no.) | 653,440 | 473,492 | 236,337 |
| Map resolution (Å) |  |  |  |
| 0.143 FSC threshold | 2.58 | 2.50 | 4.04 Å |
| Map resolution range (Å) | 2.29-6.58 | 2.20-5.11 | 4.22-7.61 |
| **Refinement** |  |  |  |
| Initial model used (PDB code) | *Ab initio* | *Ab initio* | *Ab initio* |
| Model resolution (Å) | 2.58 | 2.50 | 4.04 Å |
| FSC threshold | 0.143 | 0.143 | 0.143 |
| Model resolution range (Å) | Not applicable | Not applicable | Not applicable |
| Map sharpening B factor (Å<sup>2</sup>) | 78.3 | 83.5 | 199.1 |
| Model composition |  |  |  |
| Non-hydrogen atoms | 11948 | 11692 |  |
| Protein residues | 816 | 728 |  |
| Nucleotides | 275 | 292 |  |
| B factors (Å<sup>2</sup>) |  |  |  |
| Protein | 107 | 36 |  |
| Nucleotide | 127 | 69 |  |
| R.m.s. deviations |  |  |  |
| Bond lengths (Å) | 0.004 | 0.004 |  |
| Bond angles (°) | 0.578 | 0.604 |  |
| Validation |  |  |  |
| MolProbity score | 1.27 | 1.27 |  |
| Clashscore |  |  |  |
| Poor rotamers (%) | 0.62 | 0.84 |  |
| Ramachandran plot |  |  |  |
| Favored (%) | 98.49 | 99.02 |  |
| Allowed (%) | 1.51 | 0.98 |  |
| Disallowed (%) | 0 | 0 |  |

**Fig. 1.**
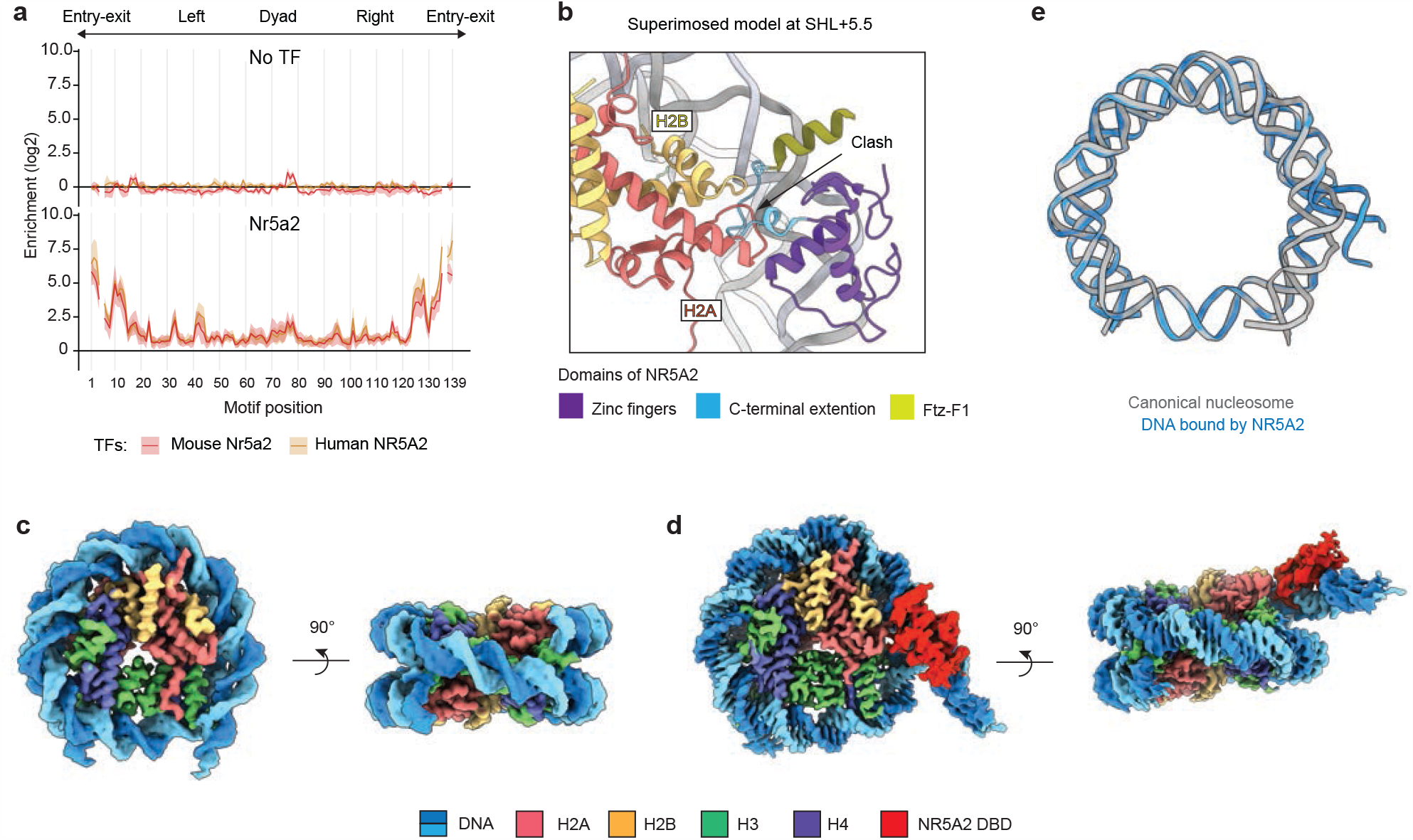
The cryo-EM structure of the human NR5A2-nucleosome complex^SHL+5.5^. (**a**) SeEN-seq enrichment profiles of mouse Nr5a2 (red) and human NR5A2 (yellow). The enrichments (log_2_) were plotted against the positions where the NR5A2 motif (TCAAGGCCA) starts along the 601 DNA. The average values of three independent experiments are shown with the SD values. Motif positions #6 and #136 were lost due to technical issues. (**b**) The superimposed model of the canonical nucleosome bound by NR5A2 DNA binding domain (PDBID: 2A66) at SHL+5.5. The CTE loop shows steric clash with histone H2A. (**c**) The cryo-EM structure of 601 nucleosome containing Nr5a2 motif at SHL+5.5. (**d**) The cryo-EM structure of human NR5A2-nucleosome complex^SHL+5.5^ at 2.58Å resolution. (**e**) Structural comparison of DNA path between canonical nucleosome and NR5A2-nucleosome complex^SHL+5.5^. The angle of DNA reorientation was measured by Pymol.

To determine which motif position to investigate further, we used modelling to superimpose the DBD onto the canonical nucleosome at the individual binding sites that had been identified by SeEN-seq (Fig. 1a). The DBD showed a good fit onto the nucleosome with no steric hindrances for motif positions at superhelical locations (SHL) -6, -5, -4, and -3. In contrast, the DBD showed clashes with histones at SHL +0.5, +2.5, +3, and +5.5 (Fig.1b and Extended Data Fig.2i,j). Based on this, we hypothesized that NR5A2 binding to these positions would affect nucleosome architecture and provide insights into the pioneer factor mechanism.

To test this, we selected the strong binding site with motif position at SHL +5.5 (Extended Data Fig. 2i). To exclude that insertion of the NR5A2 motif into 601 DNA affects the DNA end flexibility, we determined the structure of the NR5A2-unbound nucleosome^SHL+5.5^ by cryo-EM and found that it corresponds to the canonical nucleosome structure (Fig. 1c, Extended Data Figs. 3,4, and supplementary Table 1). Therefore, any structural changes in the NR5A2-bound nucleosome must be due to NR5A2 binding and not the motif insertion.

**Fig. 2.**
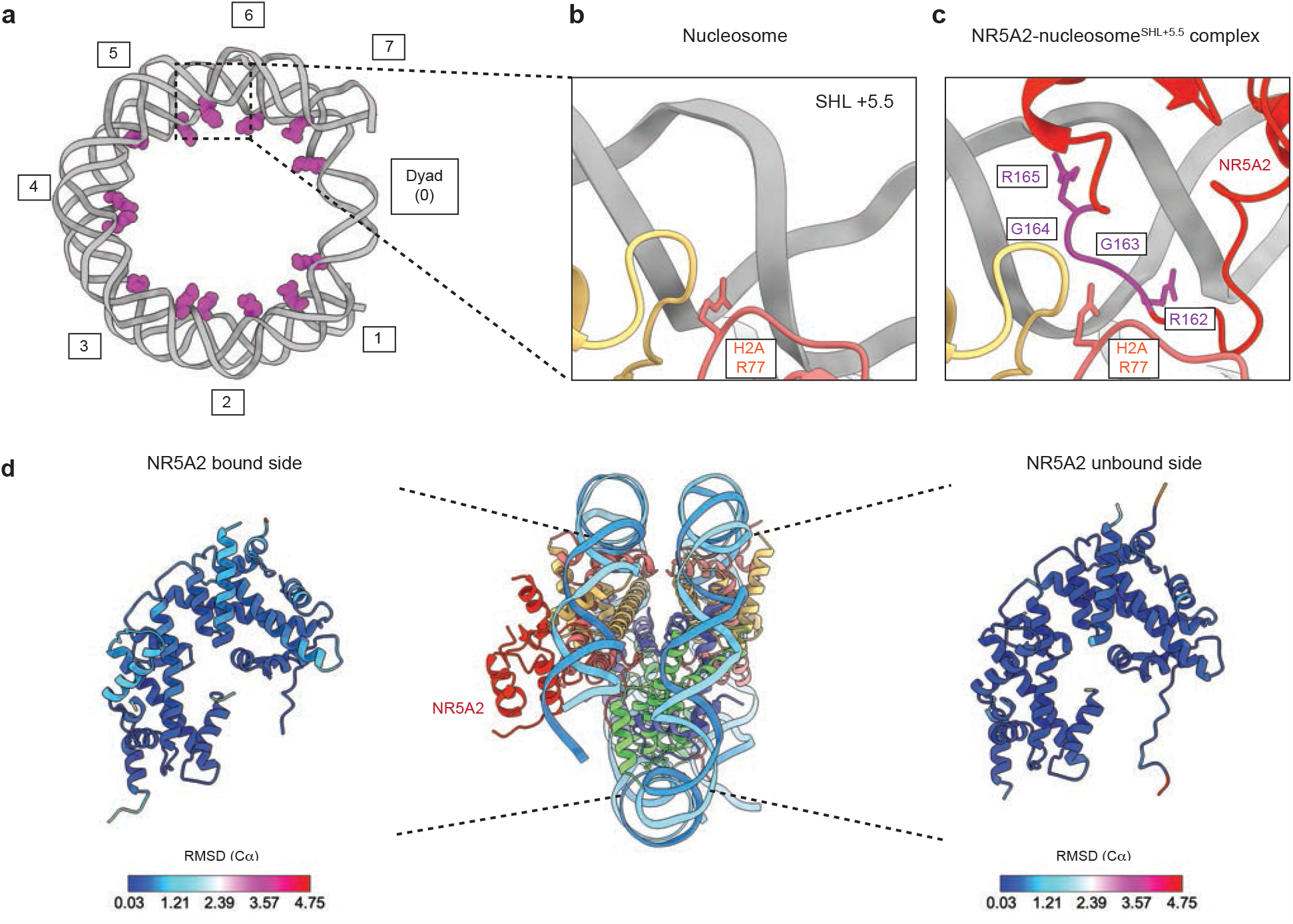
RGGR motif of the CTE loop competes with the minor groove anchor. (**a**) The nucleosome structure (PDBID: 1KX5) showing Arg (magenta) residues interacted with minor groove. Numbers indicate super helical locations. (**b**) Close-up view showing minor groove anchor H2A Arg77 at SHL+5.5 in the unbound structure. (**c**) Close-up view showing NR5A2 binding at minor groove at SHL+5.5 in the bound structure. RGGR motif (magenta) of the CTE loop occupies SHL+5.5 instead of H2A R77. (**d**) RMSD (Cα) of histones between unbound and NR5A2-bound nucleosome structure.

The structure of the human NR5A2-nucleosome complex^SHL+5.5^ was determined at 2.58 Å resolution (Fig 1d, and Extended Data Figs.3,4). It showed an extra density on the nucleosomal DNA that could be fitted well with the crystal structure of the NR5A2 DBD-DNA complex (Extended Data Fig. S4j,k). The other regions including the LBD were not visible, presumably because of the flexible nature of the hinge region (Fig. 1d). A prominent feature of the NR5A2-nucleosome complex^SHL+5.5^ is a reorientation of the DNA by ∼50° compared to the DNA path on the canonical nucleosome, demonstrating that NR5A2 binding leads to unwrapping of DNA from the histones (Fig. 1e).

### NR5A2 CTE loop competes with minor groove anchors

The nucleosome structure is stabilized by minor groove anchors, which are electrostatic interactions between arginine residues of histones and the DNA minor groove (Fig. 2a) ^26,27^. In the canonical nucleosome, the minor groove at SHL +5.5 is occupied by minor groove anchor H2A Arg77 (Fig. 2b). In contrast, the structure of the NR5A2-nucleosome complex^SHL+5.5^ shows that the minor groove is occupied by the RGGR motif of the CTE loop (Fig. 2c), suggesting that the minor groove anchor formed by H2A is outcompeted. Whether the loss of this interaction would affect nucleosome binding of NR5A2 is unclear since the major groove can still be bound by the zinc-finger domain. To test whether the minor groove interaction affects nucleosome binding, we generated a mutant NR5A2 protein in which Arg162 (RGG<u>R</u>) is replaced with Ala (R162A) (Extended Data Fig. S5a). EMSA of NR5A2 R162A showed slightly reduced DNA and nucleosome binding activities compared to wild-type protein, implying that minor groove interactions contribute to nucleosome binding (Extended Data Fig. S5b.c). However, we cannot exclude that the decrease in nucleosome binding is due to a lower DNA binding activity.

We also asked whether NR5A2 binding affects nucleosome architecture. To compare the backbone geometries of all histones, the NR5A2-nucleosome complex^SHL+5.5^ was superimposed on the nucleosome^SHL+5.5^. The root mean square deviation (RMSD) values of the corresponding Cα atoms showed no change on the NR5A2-unbound side; however, the backbone geometries of histones on the NR5A2-bound side were shifted (Fig. 2d). This suggests that histones are slightly rearranged due to NR5A2 binding, which could be due to loss of the minor groove anchor and release of DNA from histones.

### NR5A2 D159 is important for nucleosome binding

We next considered whether NR5A2 binding causes structural changes of core histones. The NR5A2-nucleosome complex showed that the side chain of H2A Lys75 (loop 2) was flipped and formed a new hydrogen bond with H2A His82 (Fig. 3a). The flipping is potentially due to a steric hindrance of H2A Lys75 with NR5A2 Asp159 since these residues collide in a superimposed model of the NR5A2-bound complex on a canonical nucleosome (Fig. 3b). The altered interactions are accompanied by a shift of H2A α2 helix towards the nucleosome center to accommodate NR5A2 binding (Fig. 3b).

**Fig. 3.**
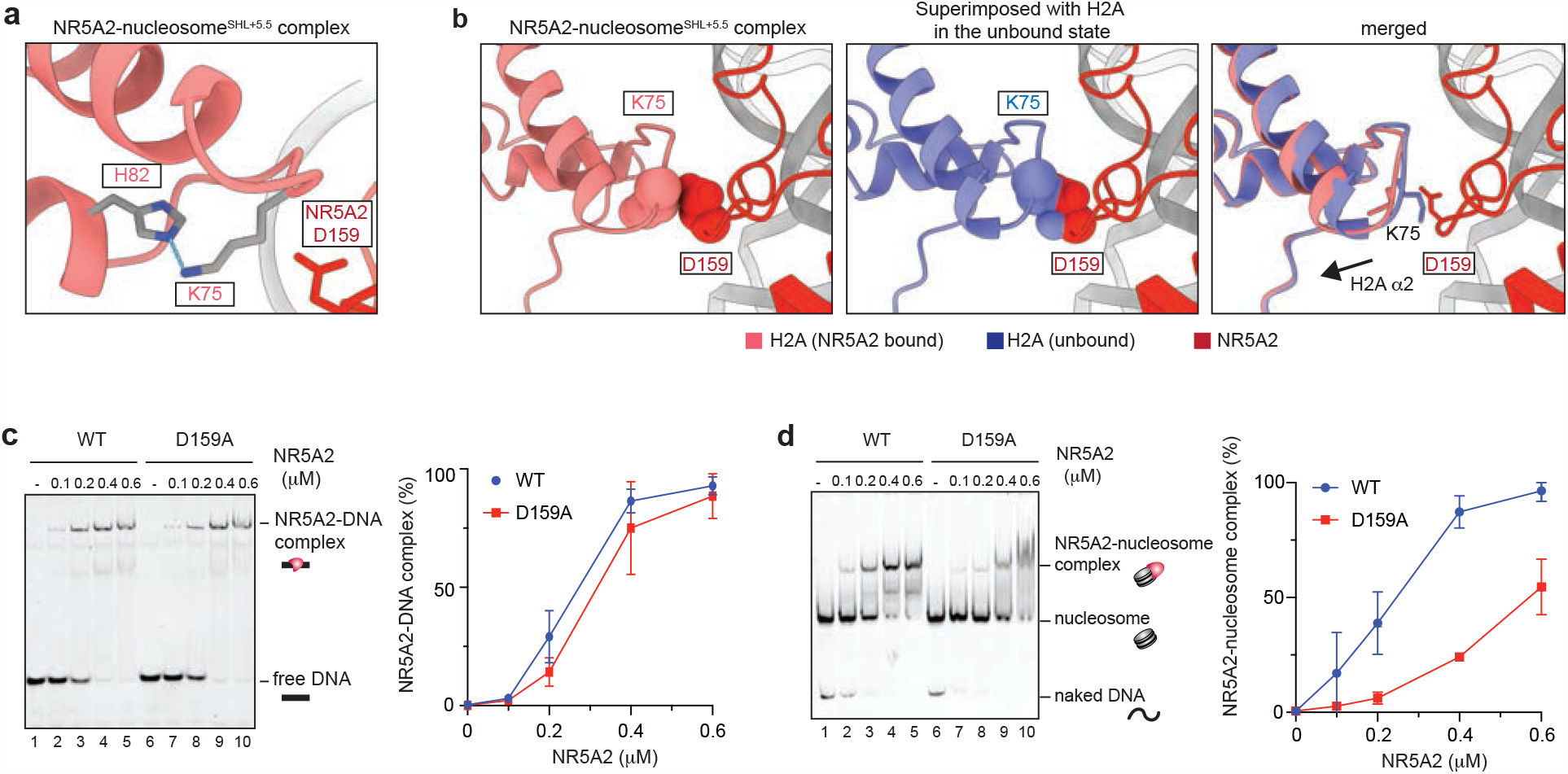
Histone rearrangements by NR5A2. (**a**) Close-up view showing the hydrogen bond between H2A K75 and H2A H82. The blue dots line indicates the distance; 2.97Å. (**b**) Close-up view showing the structural change of histone H2A induced by NR5A2 binding. Left and center; the atoms of H2A D159 and H2A K75 are shown as spheres. Right; merged model showing the rearrangement of histone H2A (**c, d**) The left panels show representative data of EMSA with naked DNA or nucleosomes containing Nr5a2 motif at SHL+5.5. Human NR5A2 wildtype (lanes 1-5) and NR5A2 D159A (lanes 6-10) were used for the experiments. Right panels show the graphical representations. The average values of three independent experiments are shown with the SD values.

To test whether NR5A2 Asp159 is required for nucleosome binding, we generated mutant NR5A2 D159A protein (Extended Data Fig. S5d). Since a defect in DNA binding would preclude further study of nucleosome binding, we first performed EMSAs with naked DNA. Mutant and wild-type protein showed no statistically significant differences in DNA binding activity (Fig. 3c). In contrast, NR5A2 D159A showed reduced nucleosome binding activity compared with wildtype (Fig. 3d). We conclude that NR5A2 Asp159 is largely dispensable for naked DNA binding but is required for nucleosome binding.

A potential limitation of these experiments is that they are based on a 601 nucleosome-positioning sequence with a motif position relatively near the DNA entry-exit sites. Whether NR5A2 can bind to its motif closer to the dyad axis on an endogenous DNA sequence and whether Asp159 would be important for this is not known. To address these questions, we selected a mouse DNA sequence that is known to be bound by Nr5a2 during ZGA ^20^ (Fig. 4a). NR5A2 wild-type and D159A mutant proteins showed similar binding affinities to the endogenous DNA sequence (Fig. 4b). Nucleosomes were reconstituted with a 149 bp sequence that contains the Nr5a2 motif in approximately SHL +2.5 (Fig. 4a). The H2A-H2B complexes used for reconstitution were labeled with Alexa647 to detect nucleosome binding (Fig. 4c). While NR5A2 efficiently bound to the nucleosome, NR5A2 D159A showed decreased binding (Fig. 4d). Therefore, NR5A2 Asp159 is also required for binding to different motif positions on a nucleosome reconstituted with an endogenous target sequence.

**Fig. 4.**
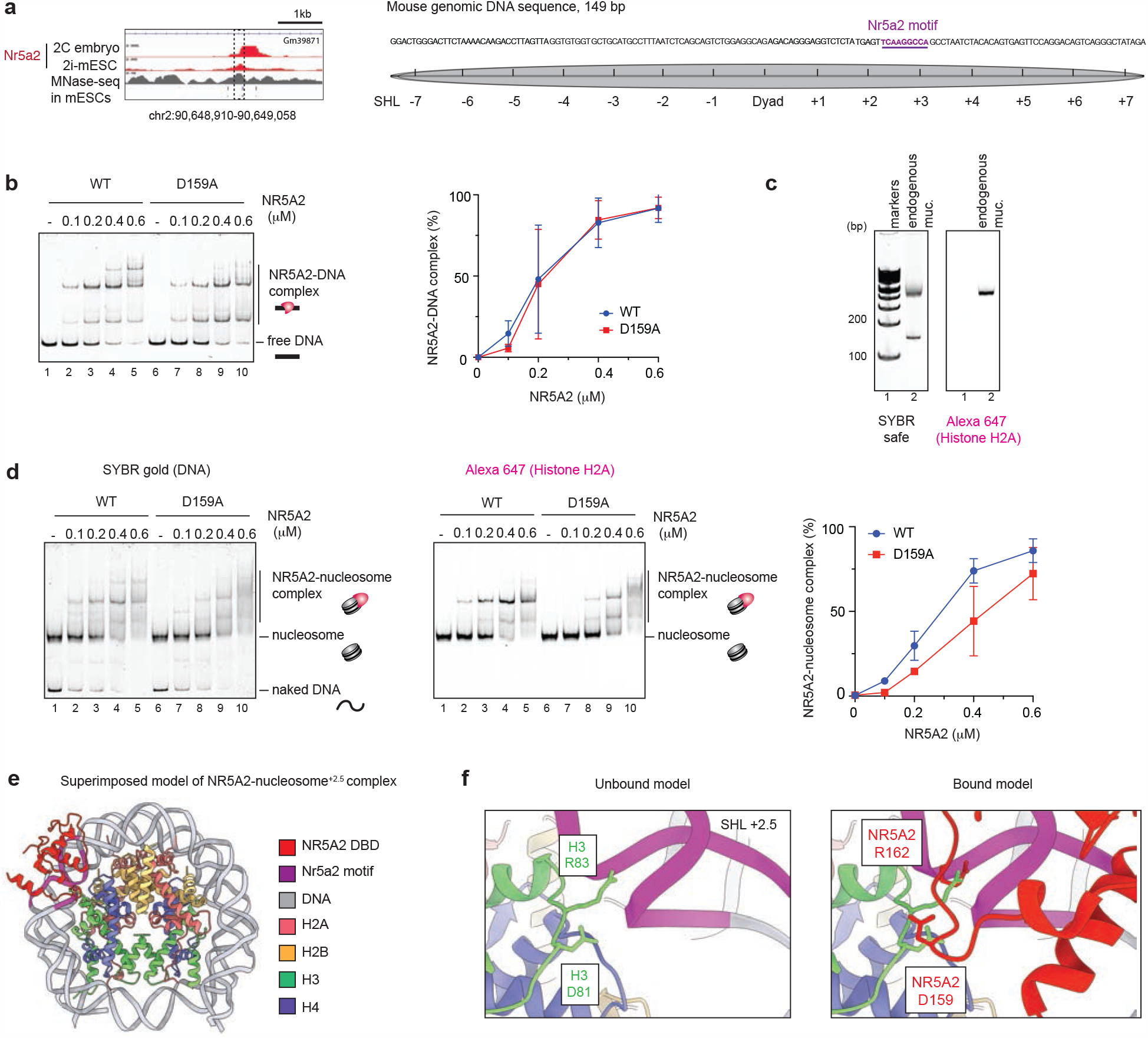
The nucleosome binding with endogenous mouse genomic nucleosome. (**a**) Left: IGV snap snapshot showing the Nr5a2 binding ^20,34^ overlapped with MNase-seq profile in mouse embryonic stem cells ^35^. The dashed rectangles highlight the region used to reconstitute mononucleosomes. Right: the DNA sequence of a 149 bp of mouse endogenous sequence used for nucleosome reconstitution. Grey circles showing possible nucleosome positions. Nr5a2 motif that locates at SHL +2.5 is shown in magenta. (**b**) The left panels show representative data of EMSA with endogenous naked DNA. The lanes 1-5 and lanes 6-10 show results with NR5A2 wildtype and NR5A2 D159A mutant, respectively. Right panels show the graphical representations. The average values of three independent experiments are shown with the SD values. (**c**) Purified endogenous mouse genomic nucleosome. Nucleosomes (lane 2) were analyzed by 6% native-PAGE and detected by SYBR gold staining and Alexa 647 fluorescence. Lane 1 indicates 100 bp markers. (**d**) The left and center panels show representative data of EMSA with the endogenous nucleosome. Alexa647 fluorescence signals correspond to the nucleosome. The lanes 1-5 and lanes 6-10 show results with NR5A2 wildtype and NR5A2 D159A mutant, respectively. Right panel shows the graphical representations. The average values of three independent experiments are shown with the SD values. (**e**) The superimposed model structure of NR5A2-nucleosome complex^SHL+2.5^. (**f**) Left: Close-up view showing minor groove anchor H3 Arg83 at SHL+2.5 in the unbound model structure. Right: Close-up view showing NR5A2 binding at minor groove at SHL+2.5 in the bound model structure. Asp159 and Arg162 in the CTE loop show steric clash with histone H3 Arg83 and Asp81, respectively.

We attempted to obtain a cryo-EM structure of this nucleosome bound by NR5A2, but this was unsuccessful due to technical reasons. We therefore built a superimposed model of NR5A2 DBD-nucleosome complex^SHL+2.5^ (Fig. 4e). NR5A2 DBD can recognize its own motif on nucleosomal DNA at SHL +2.5 in the vicinity of histones H3 and H4. The modeled structure showed that NR5A2 Arg162 in the loop region occupies the minor groove at SHL +2.5, which is bound by minor groove anchor H3 R83 (Fig. 4f). This suggests that minor groove anchor competition is also relevant for nucleosome binding when the motif is in other superhelical locations. Interestingly, NR5A2 D159 showed a clash with H3 Asp81, suggesting that H3 needs to be accommodated to allow NR5A2 binding (Fig. 4f). Given that most genomic DNA sequences bind to histone octamers less tightly than 601 DNA ^28,29^, the natural flexibility of DNA may allow efficient nucleosome binding by NR5A2. These results suggest that NR5A2 Asp159 contributes to nucleosome binding at several superhelical locations and promotes histone rearrangement.

### Dynamics of DNA unwrapping by NR5A2

The NR5A2-nucleosome structure shows that NR5A binding causes unwrapping of DNA from histones. Whether NR5A2 Asp159 promotes the release of DNA cannot be assessed using EMSA (Fig. 3e). To address this, we performed Single-molecule Förster Resonance Energy Transfer (smFRET) measures to observe the real-time dynamics of DNA unwrapping ^30-32^. To monitor the change in FRET efficiency, the donor fluorophore Cy3 was conjugated to the 5’-end of the 601 DNA, adjacent to NR5A2 motif at SHL +5.5, and the acceptor fluorophore Alexa 647 was conjugated to histone H2A K119C (Fig. 5a and Extended Data Fig.6a). Nucleosomes were reconstituted by combining Alexa647-labeled H2A-H2B histones together with H3-H4 histones, and Cy3-labeled DNA (Extended Data Fig.6b). Nucleosomes were surface-immobilized for imaging via a biotin-streptavidin interaction (Fig.5a). NR5A2 wild-type and D159A showed similar binding affinity as the nucleosome without linker DNA, suggesting that extended linker DNA does not affect nucleosome binding (Extended Data Fig.6c).

**Fig. 5.**
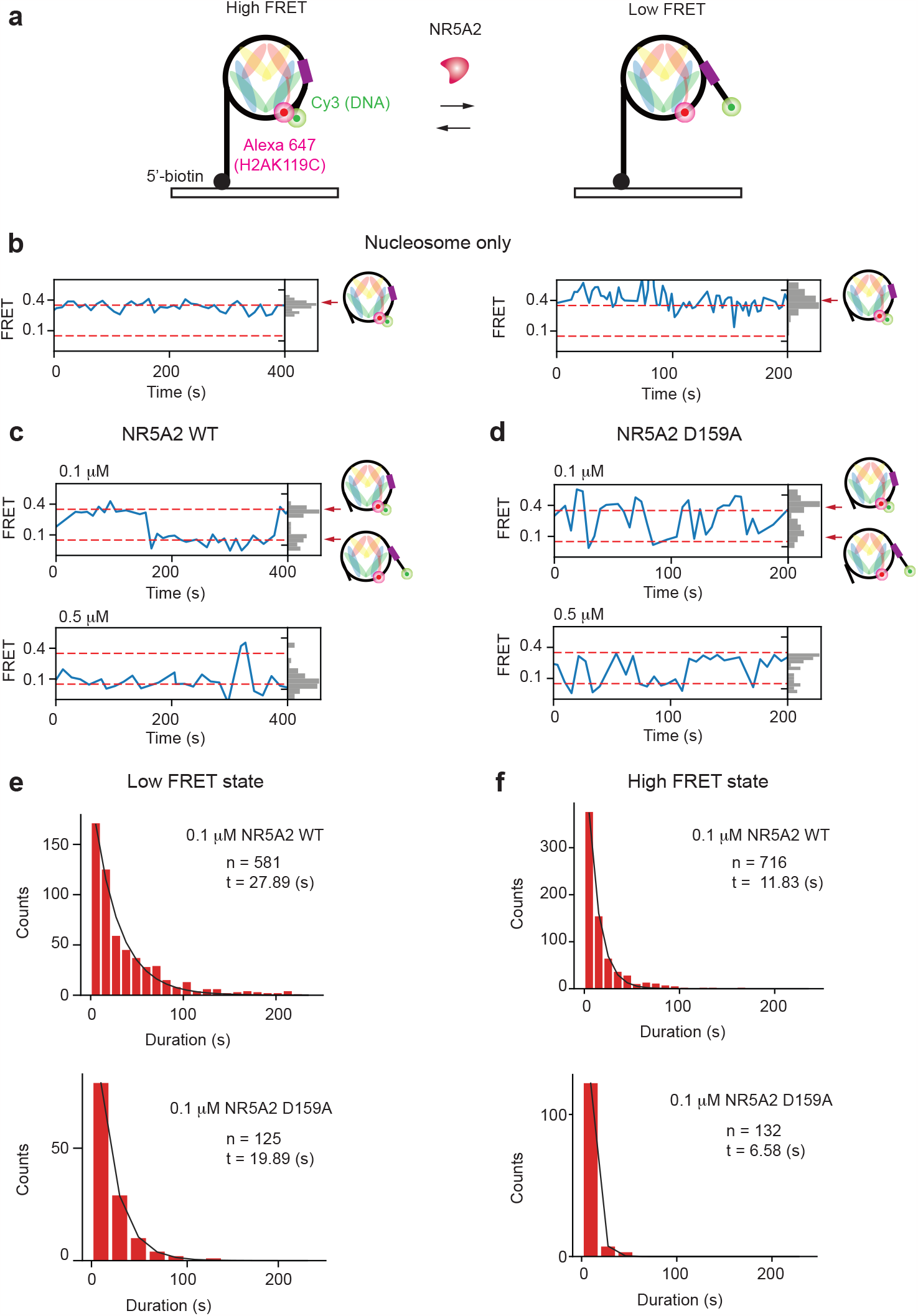
smFRET shows the dynamics of DNA unwrapping and rewrapping. (**a**) A schematic illustration showing smFRET analysis. To monitor the change in FRET efficiency, the donor fluorophore Cy3 was conjugated to the 5’-end of the 601 DNA, adjacent to Nr5a2 motif at SHL +5.5, and the acceptor fluorophore Alexa 647 was conjugated to histone H2A K119C. (**b**) Representative data of time traces of single nucleosomes. Two dot lines indicate high FRET and low FRET, respectively. (**c, d**) Representative data of time traces of single nucleosomes in the presence of NR5A2 wildtype (panel c) and NR5A2 D159A mutants (panel d). The experiments with different concentrations (0.1 μM and 0.5 μM) are shown. (**e**) Histograms showing the duration in the low FRET state with the number of counts. Upper and lower panels show the result of wildtype and D159A mutant, respectively. (**f**) Histograms showing the duration in the high FRET state with the number of counts. Upper and lower panels show the result of wildtype and D159A mutant, respectively.

In the absence of NR5A2, nucleosomes displayed a constant, high FRET state (Fig. 5b), indicating that the DNA was stably wrapped around the histone core on the timescales of these measurements. Upon addition of NR5A2 to nucleosomes, an alternation between two FRET states was observed, consistent with DNA unwrapping followed by rewrapping. Multiple transitions between high and low FRET states were observed for individual molecules (Fig. 5c, d). The dynamics observed in the presence of NR5A2 were further characterized to determine the kinetics. The NR5A2 D159A mutant displayed more frequent switching events and the duration of the low FRET state (unwrapped) was on average shorter compared to wild-type NR5A2 (NR5A2 wild-type: 27.89 s, NR5A2 D159A: 19.89 s) (Fig. 5e and Extended Data Fig.6d). This result suggests that DNA unwrapping by NR5A2 D159A is less stable compared with wild-type. Unexpectedly, NR5A2 D159A also exhibited a shorter duration of high FRET states (wrapped) (NR5A2 wild-type: 11.83 s, NR5A2 D159A: 6.58 s) before transitioning to low FRET states (Fig. 5f and Extended Data Fig.6e). Although the simplest interpretation of the FRET assay is that a low FRET state corresponds to NR5A2-bound nucleosome and a high FRET corresponds to unbound nucleosome, this cannot fully explain the shorter duration of high FRET states for the NR5A2 D159A mutant. Either this mutant binds to the nucleosome faster than wild-type or a high FRET state can also be achieved with mutant protein bound to the nucleosome. Current data do not allow us to distinguish between these possibilities. We speculate that opening and closing of the 3’ DNA entry-exit site can occur on nucleosome-bound NR5A2 D159A where mutation of the CTE prevents a stable unwrapping and the intact zinc fingers mediate DNA binding (Fig. 6). We therefore propose that CTE-mediated minor groove competition promotes a stable unwrapped state of the nucleosome.

**Fig. 6.**
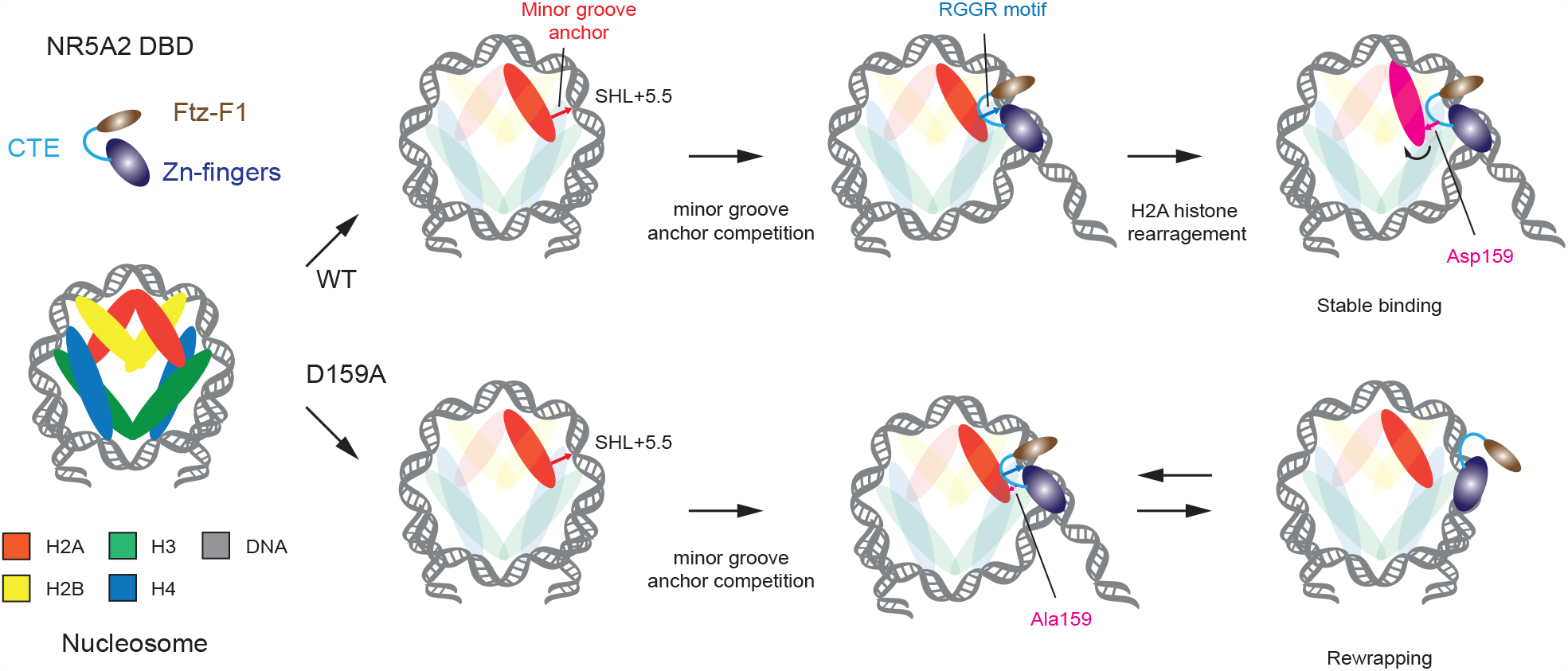
A model of DNA unwrapping by NR5A2. Zinc fingers and CTE recognize major and minor groove in the Nr5a2 motif, respectively. In the wild-type, NR5A2 weakens minor groove-histone interaction and substitutes them with the RGGR loop in CTE. CTE also induces the histone arrangement to facilitate stable nucleosome binding. Thus, CTE-mediated minor groove competition promotes a stable unwrapped state. In contrast, the CTE in D159A mutant does not stably interact with the minor groove due to the lack of histone rearrangement. The DNA binding ability is presumably retained by the zinc fingers but DNA unwrapping and rewrapping occur dynamically.

## Discussion

The structure of the NR5A2-nucleosome complex revealed a novel mechanism of minor groove anchor competition by which pioneer factors can destabilize histone-DNA interactions. Specifically, the NR5A2-nucleosome structure shows that the CTE has two functions: 1) replacement of minor groove anchors from histones, and 2) steric clashes that alter the histone arrangement and facilitate stable nucleosome binding of NR5A2 (Fig. 6). Current data do not allow us to exclude that the stable binding of NR5A2 itself is also required for DNA unwrapping. We propose that the release of DNA from the nucleosome is initiated through the minor groove anchor competition and re-wrapping of the DNA is prevented by stable binding through histone rearrangement.

The diversity in pioneer factor mechanisms is just beginning to emerge. OCT4-SOX2 and Sox11/Sox2 cause local DNA distortions and unwrap DNA when bound near the entry-exit sites ^9,10^. OCT4 uses two DNA binding domains to prevent the re-wrapping of DNA ^11^. p53 binds to linker DNA and interacts with the histone H3 tail ^33^. These pioneer factors can be classified into those that cause DNA unwrapping either by DNA distortion or wedging. Distinct from the results of these studies, our structure shows that NR5A2 weakens minor groove-histone interactions and substitutes these with the RGGR motif in the CTE loop. The exchange in minor groove interactions occurring in the NR5A2-bound nucleosome provides the first experimental evidence for a theoretical model that proposed a compensation of histone-DNA interactions with pioneer factor-DNA interactions ^35^. The loss of histone-DNA minor groove interactions is compensated by the high affinity binding of NR5A2 to DNA and results in an energetically stable nucleosome invaded by NR5A2. We note that the binding mode depends on the orientation of the motif on the nucleosome. For example, a superimposed model of the DBD on the nucleosome suggests that the CTE loop does not face the histones when the motif occurs in the same orientation on the other DNA gyre (SHL -7 to SHL 0). On the other hand, structural and modelling data suggest that the CTE loop can affect minor groove anchors at multiple positions between SHL +2.5 and +5.5 (Figs. 3 and 4). Therefore, motif position and orientation contribute to the mechanism of minor groove anchor competition by pioneer factors.

The zinc finger domain recognizes its DNA motifs via a short anchoring α-helix on the major groove, which is a common structural feature associated with pioneer factors ^13^. The NR5A2-nucleosome structure shows that the zinc finger domain binds to the major groove of DNA and recognizes the “AGGCCA” sequence on the nucleosomal DNA. The same superhelical location (SHL +5.5) bound by GATA3 as for NR5A2 shows no effect on nucleosome architecture ^12^, suggesting that major groove binding is insufficient for DNA release. The RGGR motif in the CTE targets the “TCA” sequence in the minor groove DNA. This motif is a common feature of DNA sequences that are recognized by members of the NR5A and NR3B orphan nuclear receptor families, suggesting that CTE-loop mediated minor groove competition may be conserved amongst these transcription factors ^23^ (Extended Fig. 1c). Consistent with this notion, the NR3B family member Esrrb targets its own motif on the nucleosome as a pioneer factor ^20^. A conserved mechanism of minor groove anchor competition by orphan nuclear receptors may explain their roles in directing cellular potency in natural and induced reprogramming.

## Acknowledgements

We are very grateful to Kenichiro Abe and Laura Gomez Hernandez for their contributions. We thank Katrin Straßer and Chiaki Kobayashi for technical assistance. We thank Nicolas Thomä for advice on SeEN-seq. We thank Maciej Zaczek for an initial experiment and all members of K.T.’s laboratory for discussions. We thank Tillman Schäfer at the cryo-EM facility for assistance in cryo-EM data collection and Rin Ho Kim for sequencing at the NGS facility, MPIB.

## Funding

European Research Council grant ERC-CoG-818556 TotipotentZygotChrom (KT) Max Planck Society (KT)

## Author contributions

WK and RH performed protein purification and sample preparation for cryo-EM analysis. MK performed SeEN-seq. SR analyzed SeEN-seq data. WK, DB, and JB performed cryo-EM analysis and model building. WK and EA performed biochemical experiments. AS and KD performed smFRET analyses. KT conceived the project and supervised the work. WK, AS, KD, and KT planned the project, designed the experiments, and wrote the manuscript. All authors discussed the results and commented on the manuscript.

## Competing interests

The authors declare that they have no competing interests.

## Data and materials availability

Requests for plasmids generated in this study should be directed to the corresponding author. The dataset of Nr5a2 CUT&Tag and ChIP-seq were obtained from GSE178234 and GSE92412, respectively. MNase-seq data in mESCs was obtained from GSE82127.

## Notes

### Competing Interest Statement

The authors have declared no competing interest.

