## Supplementary figures for "Nucleosome-bound NR5A2 structure reveals pioneer factor mechanism by minor groove anchor competition"

#### Online methods

##### Protein purification

$His_6$-tagged full-length mouse Nr5a2 was purified as described previously [20]. The DNA fragment encoding human NR5A2 was ligated into NdeI-BamHI sites of pET-15b vector with an N-terminal $His_6$-TEV tag. Human NR5A2 was purified by Ni-NTA agarose (Qiagen) the same as mouse Nr5a2 purification. $His_6$-tag was removed with TEV protease during dialysis, and NR5A2 was further purified by SP sepharose fast flow (Cytiva). The purified sample was dialyzed against buffer 5 (20 mM Tris-HCl (pH 7.5), 400 mM NaCl, 5% glycerol and 1 mM DTT). Human NR5A2 R159A and NR5A2 162A mutants were purified the same as wild-type.

Mouse histones H2A, H2A K119C, H2B, H3.3, and H4 were expressed and purified as previously described [36]. Histone octamer was reconstituted as described previously [36]. Histones H2AK119C-H2B complex and H3.3-H4 complex were reconstituted as previously described. For the fluorescent labeling of H2A-H2B complex, 50 mM of H2A K119C-H2B complex were incubated with 500 mM Alexa Flour 647 $C_2$ Maleimide (Invitrogen) in 20 mM Tris-HCl (pH 7.5) buffer containing 0.1 M NaCl and 1 mM TCEP at room temperature for 2h in the dark. The reaction was stopped by the addition of 150 mM of 2-mercaptoethanol, and the sample was then dialyzed against 20 mM Tris-HCl buffer containing 0.1M NaCl, 1 mM EDTA, and 5mM 2-mercaptoethanol.

##### SeEN-seq assay

SeEN-seq was performed as previously described[10,20]. Nr5a2 motif (JASPAR ID: MA0505.1, TCAAGGCCA) was tiled 1 bp interval across 147 bp Widom 601 DNA sequence. The pooled DNA library (153 bp) was digested by EcoRV and was further purified by a gel-extraction kit. For the preparation of the nucleosome library, purified library DNA (3 μg) was mixed with spike-in 153 bp 601 DNA (97 μg). The nucleosome library was reconstituted by salt dialysis method and was further purified by polyacrylamide gel (6%) electrophoresis using a Prep Cell apparatus (Bio-Rad). The nucleosomes (0.1 μM) were incubated with mouse Nr5a2 (0, 1, or 1.2 μM) or human NR5A2 (0, 0.4, 0.8 μM) at room temperature for 30 min in a reaction buffer (20 mM Tris-HCl (pH7.5), 120 mM NaCl, 1 mM $MgCl_2$, 10 μM $ZnCl_2$, 1 mM DTT, 100 μg/ml BSA). Three independent experiments were performed, and the bands were visualized by SYBR Gold staining (Invitrogen). DNA libraries were prepared as described previously. Purified DNA libraries were sequenced on a NextSeq 500 with paired-end 150 bp reads (Illumina). The DNA sequences are shown in supplementary tables S1.

##### SeEN-seq data processing

SeEN-seq data analysis was performed as described previously[10, 20]. In brief, raw reads were mapped to the reference constructs used in SeEN-seq assay by Bowtie2 [37] (version 2.3.5.1) with the following parameters: -t -q –very-sensitive –no-discordant –no-mixed. Only reads with map quality higher than 20 were used for further analysis steps. Samtools idxstat [38] command was used to count the number of reads that map to each construct of the SeEN-seq library. The read counts were normalized by their respective library size, before further dividing by the library size normalized value of counts of the Widom 601 template construct in their library. Finally, the transcription factor binding enrichment score of each construct is represented as a $log_2$-transformed fold-change between normalized bound and unbound fraction of each construct. The raw read numbers are shown in supplementary tables S1.

##### Preparation of the nucleosomes

601 DNA The endogenous DNA fragment for nucleosome reconstitution were amplified by PCR and further purified by polyacrylamide gel (6%) electrophoresis using a Prep Cell apparatus (Bio-Rad). The eluted DNA was concentrated with an Amicon Ultra centrifugal filter unit (Millipore).

The nucleosomes were reconstituted by salt dialysis method. For the nucleosome reconstitution with 601 DNA, the DNA and mouse histone octamer were mixed in a 1:1.8-2.0 molar ratio in 2M KCl. For the nucleosome reconstitution with endogenous DNA sequence, the DNA, Alexa 647-labeled histone H2A-H2B complex, and H3.3-H4 complex were mixed in 1:4:3.6 molar ratio in 2M KCl high salt buffer. The reconstituted nucleosomes were further purified by polyacrylamide gel (6%) electrophoresis using a Prep Cell apparatus (Bio-Rad). The nucleosomes were concentrated with an Amicon Ultra centrifugal filter unit (Millipore) and were stored in TCS buffer (20 mM Tris-HCl (pH 7.5) and 1 mM DTT) at 4°C.

**Preparation of NR5A2-nucleosome complex for cryo-EM analysis**

The nucleosome (2 µM), in which Nr5a2 motif locates at 128bp from the 5' end of DNA (54 bp from the dyad, SHL +5.5), was mixed with human NR5A2 (4 µM) in a reaction buffer (20 mM Tris-HCl (pH7.5), 120 mM NaCl, 1 mM $MgCl_2$, 10 µM $ZnCl_2$, 1 mM DTT). After incubation at 25°C in the heat block for 30 min, the sample was purified and stabilized by the Grafix method [39]. A gradient was formed with buffer 1 (10 mM HEPES-NaOH (pH 7.5), 20 mM NaCl, 1 µM $ZnCl_2$, 1 mM DTT, and 5% sucrose) and buffer 2 (10 mM HEPES-NaOH (pH 7.5), 20 mM NaCl, 1 µM $ZnCl_2$, 1 mM DTT, 20% sucrose, and 2% formaldehyde), using a Gradient Master (BioComp). The reconstituted NR5A2–nucleosome complex$^{SHL+5.5}$ (150 µl x 2) was placed on the top of the gradient solution and was ultracentrifuged at 27,000 rpm at 4 °C for 16 h, using an SW41 Ti rotor (Beckman Coulter). The fractions were analyzed by 5% non-denaturing PAGE. The collected samples were then desalted with a PD-10 column (GE Healthcare), equilibrated with 10 mM HEPES-NaOH (pH 7.5) buffer containing 1 µM $ZnCl_2$, and 1 mM DTT and were concentrated with an Amicon Ultra centrifugal filter unit (Millipore). The sample of nucleosomes without NR5A2 was purified by the same method.

**Cryo-EM specimen preparation and data acquisition**

To prepare the cryo-EM specimen, the sample (4 µl) was applied to a glow-discharged holey carbon grid (Quantifoil R1.2/1.3 200-mesh Cu). The grids were blotted for 3.0 s at a blotting strength setting of "5" under 100% humidity at 4 °C and then plunged into liquid ethane, cooled by liquid nitrogen, using a Vitrobot Mark IV (Thermo Fisher). Data acquisition for the NR5A2-nucleosome complex$^{SHL+5.5}$ was conducted using a 300 kV Titan Krios G2 (Thermo Fisher Scientific) equipped with a GIF Quantum 967 energy filter and a K3 direct electron detector (Gatan) running in correlated double sampling mode at a magnification factor of 105kx, equivalent to a pixel size of 0.85 Å/px. A total of 12,510 movies were collected, with a total electron dose of 65.4 e/Å² fractionated over 40 frames. The data set for pure nucleosomes was acquired on a 200 kV Glacios (Thermo Fisher Scientific) equipped with a K2 Summit direct electron detector (Gatan) at a magnification factor of 22kx, equivalent to a pixel size of 1.89 Å/px. Here, 1,998 movies were collected, with a total electron dose of 61 e/Å² fractionated over 40 frames.

**Image processing**

Data processing was performed by CryoSPARC [40]. For the dataset obtained by Gracios, data processing was performed on-the-fly via CryoSPARC Live (version 4.1) on its default processing pipeline, applying C2 point group symmetry for the final homogeneous refinement. This resulted in a 3D coulomb potential map at a nominal resolution of 4.04 Å. This map was subsequently post-processed via local b-factor correction by DeepEMhancer [41].

**Model building and structural analysis**

We have placed a nucleosome model by molecular replacement using the PDBID 3LZ0 as template. In the case of the NR5A2-nucleosome we placed by rigid body fitting in the EM map the NR5A2 model using the PDBID 2A66. The models for the free nucleosome and NR5A2-nucleosome complex were completed by manual rebuilding in COOT [42] and refined using in Phenix real space refinement [43]. The structures were analyzed using ChimeraX [44]. Molecular models were superimposed with chain A using the matchmaker option. The RMSD calculation for each residue was calculated using ChimeraX.

**Electrophoretic mobility shift assay (EMSA)**

For EMSA with short oligo DNA, 25 bp dsDNA (50 nM) containing Nr5a2 consensus motif (GAGAGAG<u>TCAAGGCCA</u>TGGCTCACT) was incubated with NR5A2s at room temperature for 30 min in a reaction buffer (20 mM Tris-HCl (pH7.5), 120 mM NaCl, 1 mM $MgCl_2$, 10 µM $ZnCl_2$, 1 mM DTT,

100 μg/ml BSA). After the incubation, the samples were loaded onto 10% non-denaturing polyacrylamide gels (0.5xTBE), and electrophoresis was performed at 100V for 1 hour at room temperature. The gels were stained by SYBR Gold (Invitrogen) and were imaged using GelDoc Go imaging system (Bio-Rad).

For EMSA with nucleosome, the nucleosomes (50 nM) were incubated with NR5A2s at room temperature for 30 min in a reaction buffer (20 mM Tris-HCl (pH7.5), 120 mM NaCl, 1 mM $MgCl_2$, 10 μM $ZnCl_2$, 1 mM DTT, 100 μg/ml BSA). After the incubation, the samples were loaded onto 10% non-denaturing polyacrylamide gels (0.5xTBE), and electrophoresis was performed at 100V for 70 min at room temperature. The gels were imaged by detecting Alexa 647 fluorescence and SYBR Gold (Invitrogen) using ChemiDoc MP imaging system or GelDoc Go imaging system (Bio-Rad).

For quantification analysis, the data from at least 3 replicates were analyzed using ImageLab (Bio-Rad) and plotted in Prism (GraphPad). Error bars show mean and standard deviation.

**Single-molecule FRET assay**

**PEG-Biotin microscopy slide preparation**

Glass coverslips for TIRF microscopy were prepared as described previously[45]. In brief, 22x22 mm cover slips (Marienfeld) were cleaned extensively, silanized with 3- aminopropyltriethoxysilane in acetone and incubated with a PEG-/PEG-Biotin solution (0.4% (w/v) Biotin-PEGSuccinimidyl Carbonate (MW 5,000) and 15% (w/v) mPEG-Succinimidyl Carbonate (MW 5,000) in fresh 0.1 M $NaHCO_3$) over night. Afterwards, they were washed with $ddH_2O$, dried under an air stream and stored under vacuum.

**Flow cell preparation**

For flow cell assembly a glass coverslip was incubated with 0.2 mg/ml streptavidin in blocking buffer (50 mM Tris, pH 7.6, 50 mM KCl, 2 % (v/v) Tween20) for 20 minutes. It was then rinsed with $ddH_2O$ and dried under an air stream. A double-sided tape (~ 0.1 mm thick) with two flow lanes of approximately 0.5 mm width was taped to a glass slide containing entry and exit holds for buffer tubes and then to the functionalized cover slip forming two flow lanes. Polyethylene tubes of 25 cm length (0.58 mm inner diameter) were stuck into entry and exit holds of the flow lanes and immobilized using epoxy glue. The flow cell was further glued into a custom-build metal holder prohibiting air diffusion to the sample. The holder was then mounted onto a home-build TIRF microscope. The flow lane surface was coated by flowing in blocking buffer for 5 minutes at 100 μl/minute and incubation for at least 15 minutes. It was then washed with working buffer (50 mM Tris pH 7.6, 120 mM NaCl, 1 mM $MgCl_2$, 10 μM $ZnCl_2$, 10 %Glycerol, 1 mM DTT, 100 μg/ml BSA) at 100 μl/min for 5 minutes.

**Sample preparation and application**

All buffers were degassed prior to use at 40 mbar for 40 minutes. Nucleosomes were diluted in working buffer (50 mM Tris pH 7.6, 120 mM NaCl, 1 mM $MgCl_2$, 10 μM $ZnCl_2$, 10 %Glycerol, 1 mM DTT, 100 μg/ml BSA) to a concentration of about 800 fM. They were applied to the flow lane at 150 μl/min for 30 seconds, incubated on the slide for 20 seconds and washed out extensively at 150 μl/min for 5 minutes. A second wash with imaging buffer (working buffer with 1 mM Trolox (aged for 5 minutes under UV light), 2.5 mM PCA, 0.21 U $ml^{-1}$ PCD) was performed (1 minute, 150 μl/min). Nr5a2 was applied at different concentrations (either 0.1 or 0.5 μM) in imaging buffer by flowing it in for 2 to 3 minutes at 150 μl/minute. Then, the flow was stopped and the tubes clipped to avoid oxygen diffusion. Data acquisition was initiated immediately.

**Imaging conditions**

A RM21-micro-mirror TIRF microscope from Mad City Labs (MCL, Madison, Wisconsin, USA) modified according to Larson *et al.* (2014)[46] equipped with an Apo N TIRF 60 x oil immersion objective (NA 1.49, Olympus) was used. The room temperature was controlled at 22.5 ± 0.5C. Green laser pulses (OBIS 532 nm LS 120 mW, Coherent) of 200 ms duration and 2 V were used to excite Cy3 dyes on the nucleosomes. Scattered light was removed, and emission was separated with emission filter sets (ET520/40 m and ZET532/640 m, Chroma) as well as split at 635 nm using an OptoSplit II dualview (Cairn Research, UK).

It was then detected with an iXon Ultra 888 EMCCD camera (Andor). Image acquisition frequency ranged from 6 s for Nr5a2 D159A to 10 s for Nr5a2. Auto focusing was performed every 8 to 10 frames while videos of up to 30 minutes were collected. A total of up to one hour of measurements was performed on each sample before nucleosome stability decreased.

**Data analysis**

FRET data was analyzed as reported [47] using the Fiji plugin Mars (Molecule Archive Suite). All images were corrected for the gaussian beam profile of the excitation light source. Populations of nucleosomes containing only a Cy3 dye (Donor-only) and nucleosomes with both dyes, displaying FRET, were selected by finding spots in the red detector region and converting them to the green region using an Affine2D matrix. Single spots were integrated, the immediate background subtracted and the fluorescence over time analyzed for each molecule separately. Bleaching positions for both Donor and Acceptor fluorophores were determined by finding the largest change point within the signal. FRET traces were selected based on several criteria: Accepted events could only display one bleaching event per fluorophore, show a FRET signal for several frames before bleaching, as well as Donor fluorescence recovery after bleaching of the Acceptor or vice versa. To investigate the dwell times of Nr5a2 on nucleosomes only FRET traces with switching events between high and low FRET states were then chosen for further analysis. These events are indicated by a sudden increase or decrease in Donor fluorescence and an anticorrelated change in FRET signal.

The FRET Efficiency was calculated for each molecule separately after correcting the signal for leakage from green to red channel (alpha) and the differences in detection efficiencies of the two channels as well as quantum yields of the two dyes (gamma) performed as previously reported [48].

To determine residence times of Nr5a2 on the nucleosomes, changes in FRET efficiency over time were investigated. A threshold of 0.48 was chosen to separate low and high Efficiency values corresponding to Nr5a2 dwell times on nucleosomes and times with free, unbound nucleosomes. The lifetimes of the states were fitted using single exponential decay curves. Binding and unbinding rates were calculated by inverting the half-lives of the respective states.

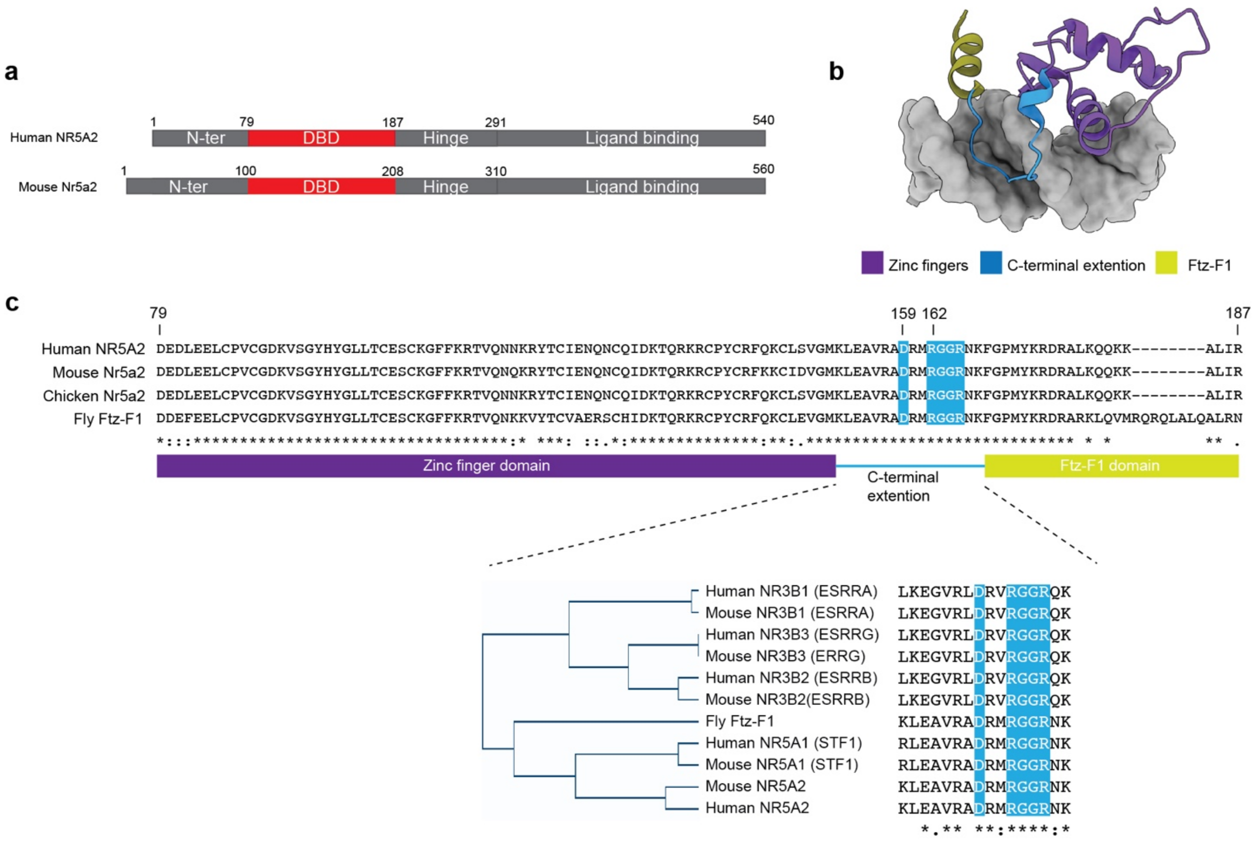


**Extended Fig. 1 Sequence alignment of Nr5a2**

(**a**) Schematic representation showing the domains of mouse and human Nr5a2. Red squares indicate the DNA-binding domain. (**b**) The crystal structure of human NR5A2-DNA complex (PDBID: 2A66). Zinc-finger domains, C-terminal extension (CTE), and Ftz-F1 domain are shown in purple, blue, and yellow, respectively. (**c**) Sequence alignment showing DNA-binding domain of Nr5a2 (human, mouse, and chicken) and fly Ftz-F1 proteins. The number of amino acids shows human NR5A2. The conservation of the CTE loop in NR5A and NR3B families is shown below.

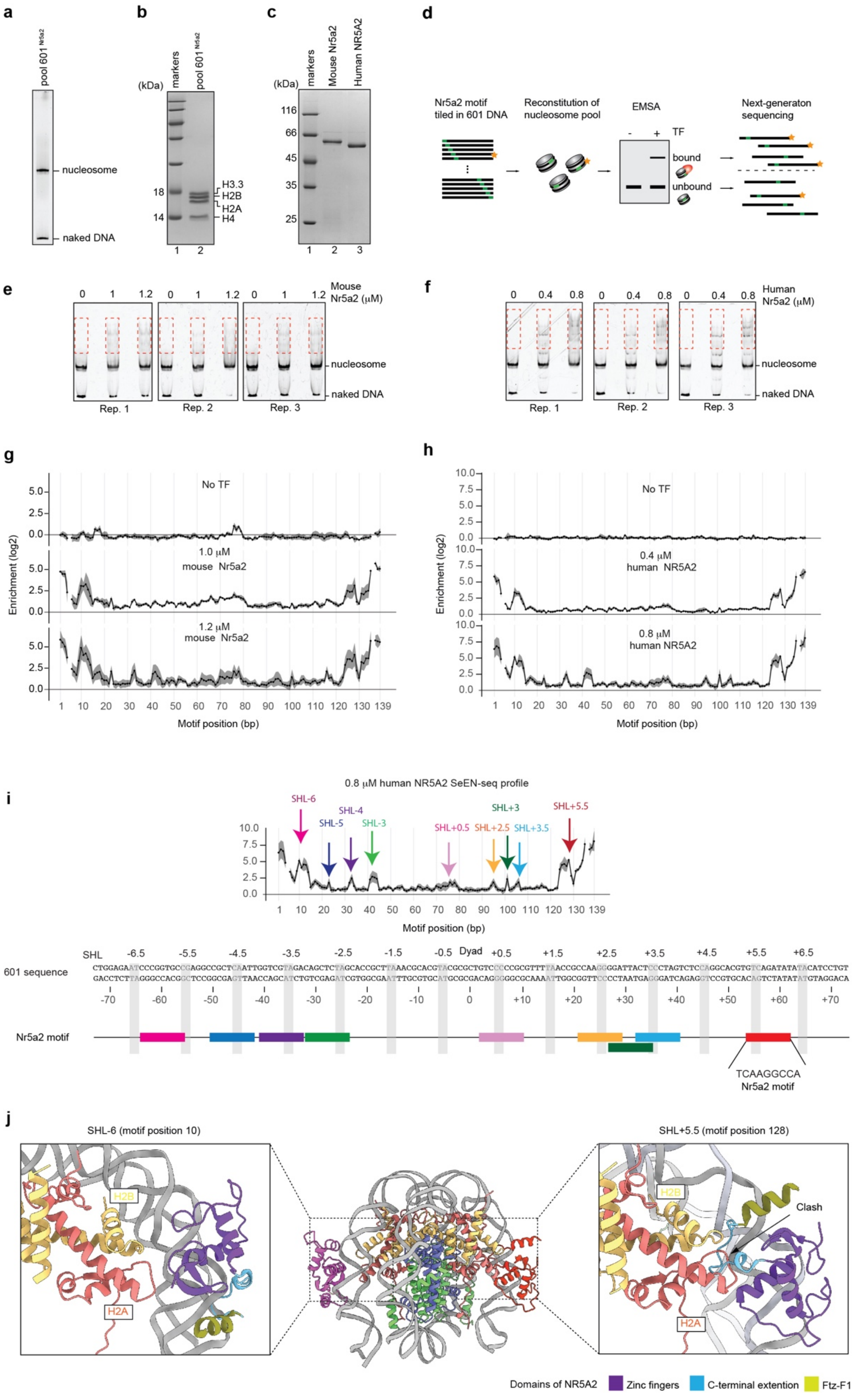
a
pool 601 Nr5a2
nucleosome
naked DNA
b
(kDa)
markers
pool 601 Nr5a2
18
14
H3.3
H2B
H2A
H4
1 2
c
(kDa)
markers
Mouse Nr5a2
Human NR5A2
116
66
45
35
25
1 2 3
d
Nr5a2 motif tiled in 601 DNA
Reconstitution of nucleosome pool
EMSA
- + TF
bound
unbound
Next-generaton sequencing
e
0 1 1.2 0 1 1.2 0 1 1.2
Mouse Nr5a2 (μM)
nucleosome
naked DNA
Rep. 1
Rep. 2
Rep. 3
f
0 0.4 0.8 0 0.4 0.8 0 0.4 0.8
Human Nr5a2 (μM)
nucleosome
naked DNA
Rep. 1
Rep. 2
Rep. 3
g
No TF
1.0 μM mouse Nr5a2
1.2 μM mouse Nr5a2
Enrichment (log2)
Motif position (bp)
h
No TF
0.4 μM human NR5A2
0.8 μM human NR5A2
Enrichment (log2)
Motif position (bp)
i
0.8 μM human NR5A2 SeEN-seq profile
SHL-6
SHL-5
SHL-4
SHL-3
SHL+0.5
SHL+2.5
SHL+3
SHL+3.5
SHL+5.5
Motif position (bp)
SHL
Dyad
601 sequence
Nr5a2 motif
TCAAGGCCA
Nr5a2 motif
j
SHL-6 (motif position 10)
SHL+5.5 (motif position 128)
H2B
H2A
Clash
Domains of NR5A2
Zinc fingers
C-terminal extention
Ftz-F1

**Extended Fig. 2 SeEN-seq analysis**

(**a**) Purified nucleosome library containing Nr5a2 motif (pool $601^{Nr5a2}$) was analyzed by native-PAGE with SYBR safe staining. (**b**) The histone contents of the purified nucleosome library were analyzed by 18% SDS-PAGE. (**c**) Purified mouse Nr5a2 (lane 2) and human NR5A2 (lane 3) used for SeEN-seq analysis. Lane1 indicates molecular markers. (**d**) Schematic illustration of SeEN-seq analysis. EMSA results were shown in panels E and F. (**e and f**) EMSA with mouse or human Nr5a2. Three independent experiments were performed. Red dot squares indicate sliced regions as bound fractions. (**g, h**) SeEN-seq enrichment profiles of mouse Nr5a2 and human NR5A2. The enrichments ($log_2$) were plotted against the positions where the NR5A2 motif (TCAAGGCCA) starts along the 601 DNA. The average values of three independent experiments are shown with the SD values. Motif positions #6 and #136 were lost due to technical issues. (**i**) Enriched sites identified by SeEN-seq analysis. Each color indicates the Nr5a2 motif in the different positions in the nucleosomal DNA. (**j**) The superimposed model of the canonical nucleosome bound by NR5A2 DBD. NR5A2 DBD does not show steric clash with histones at SHL -6 but not SHL +5.5.

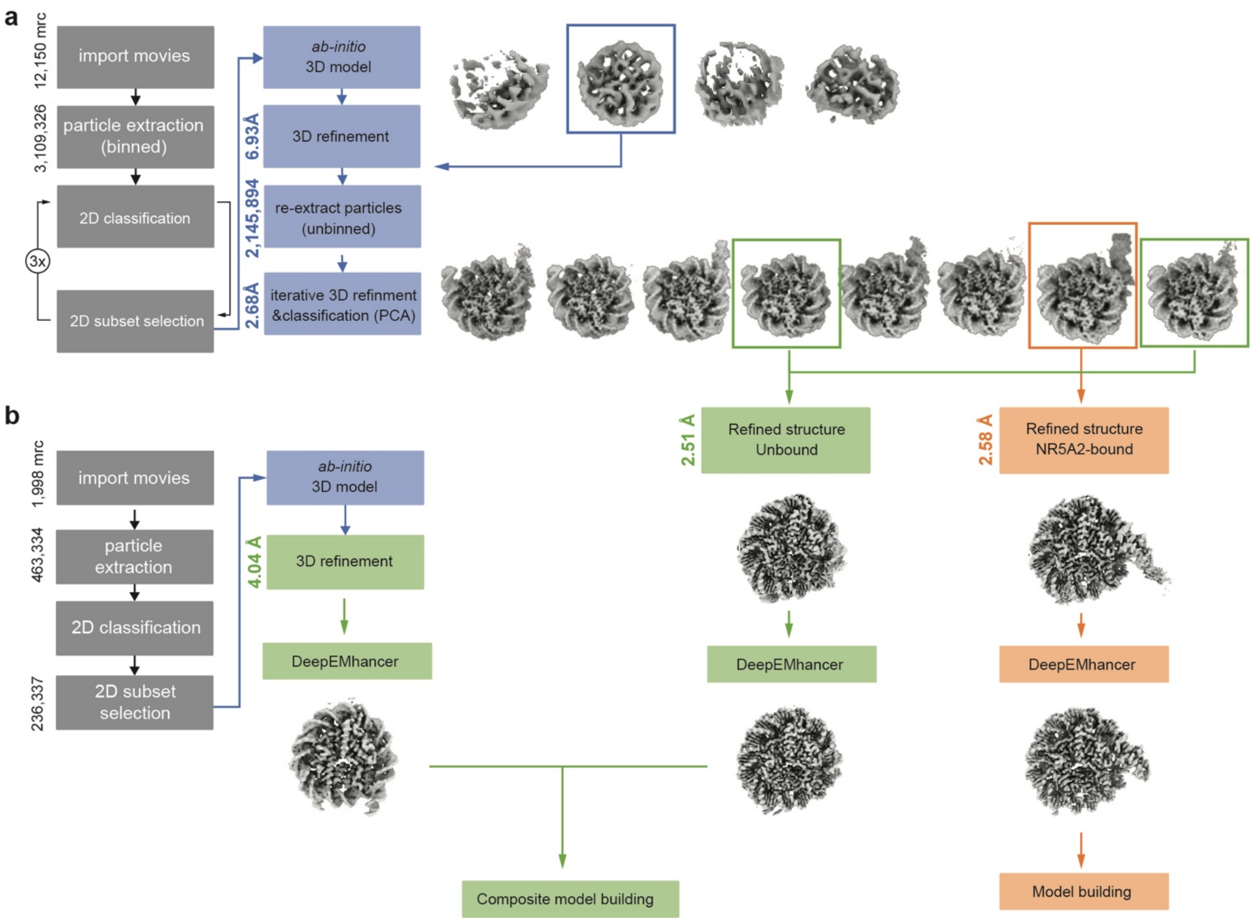


**Extended Fig. 3. Cryo-EM data processing flowchart for unbound and NR5A2-bound nucleosome**

(**a**) Flowchart of the dataset obtained by Titan Krios G2. (**b**) Flowchart of the dataset obtained by Glacios. Two EM maps were composited for the model building of unbound nucleosome.

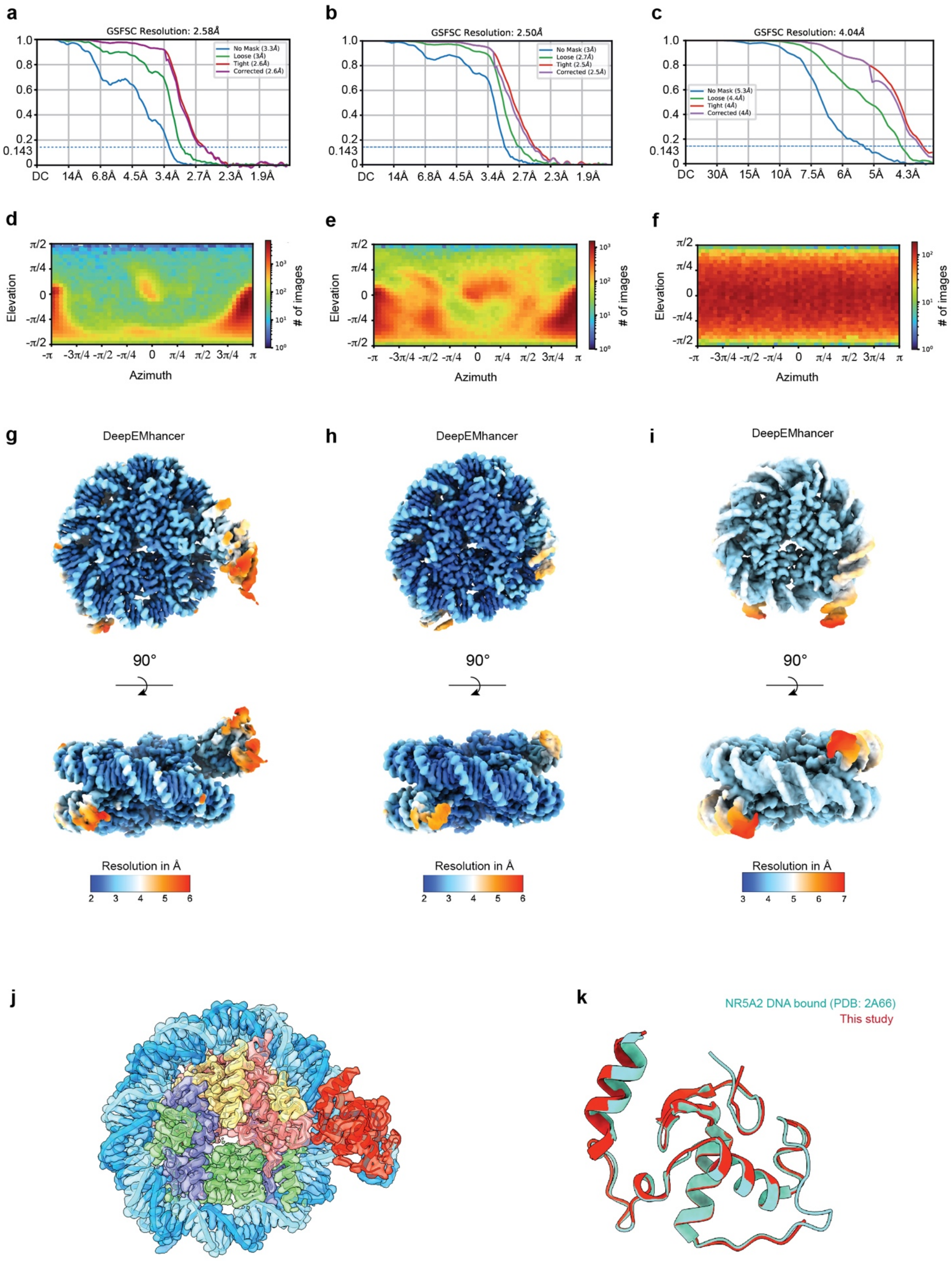

a
GSFSC Resolution: 2.58Å
No Mask (3.3Å)
Loose (3Å)
Tight (2.6Å)
Corrected (2.6Å)
b
GSFSC Resolution: 2.50Å
No Mask (3Å)
Loose (2.7Å)
Tight (2.5Å)
Corrected (2.5Å)
c
GSFSC Resolution: 4.04Å
No Mask (5.3Å)
Loose (4.4Å)
Tight (4Å)
Corrected (4Å)
d
e
f
Elevation
Azimuth
### of images
g
h
i
DeepEMhancer
90°
Resolution in Å
j
k
NR5A2 DNA bound (PDB: 2A66)
This study

**Extended Fig. 4 Data quality of NR5A2-nucleosome complex$^{SHL+5.5}$**

(**a-c**) Fourier Shell Correlation (FSC) curves of maps with the FSC = 0.143 criterion. (**d-f**) Angular distribution plot of particles employed to reconstruct maps. (**g-i**) Local resolution of maps. (**j**) Model structure fitted to the map of NR5A2-nucleosome complex$^{SHL+5.5}$. (**k**) Superimposition of the human NR5A2 DBD bound to naked DNA (PDB: 2A66) and NR5A2 DBD bound to nucleosome (this study).

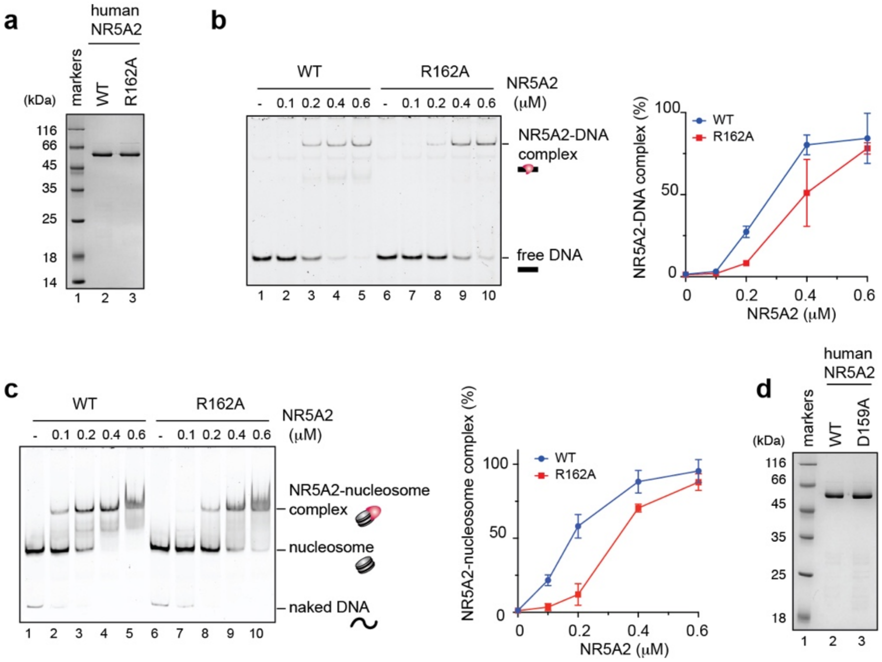


**Extended Fig. 5 Purification of NR5A2 mutants and EMSA**

(**a**) Purified human NR5A2 wildtype (lane 2) and human NR5A2 R162A mutant (lane 3) were analyzed by 12% SDS-PAGE. Lane1 indicates molecular markers. (**b**) The left panels show representative data of EMSA with naked DNA containing Nr5a2 motif. Human NR5A2 wildtype (lanes 1-5) and NR5A2 R162A (lanes 6-10) were used for the experiments. Right panels show the graphical representations. The average values of three independent experiments are shown with the SD values. (**c**) The left panels show representative data of EMSA with 601 nucleosome containing Nr5a2 motif at SHL+5.5. Human NR5A2 wildtype (lanes 1-5) and NR5A2 R162A (lanes 6-10) were used for the experiments. Right panels show the graphical representations. The average values of three independent experiments are shown with the SD values (**d**) Purified human NR5A2 D159A mutant (lane 3) were analyzed by 12% SDS-PAGE.

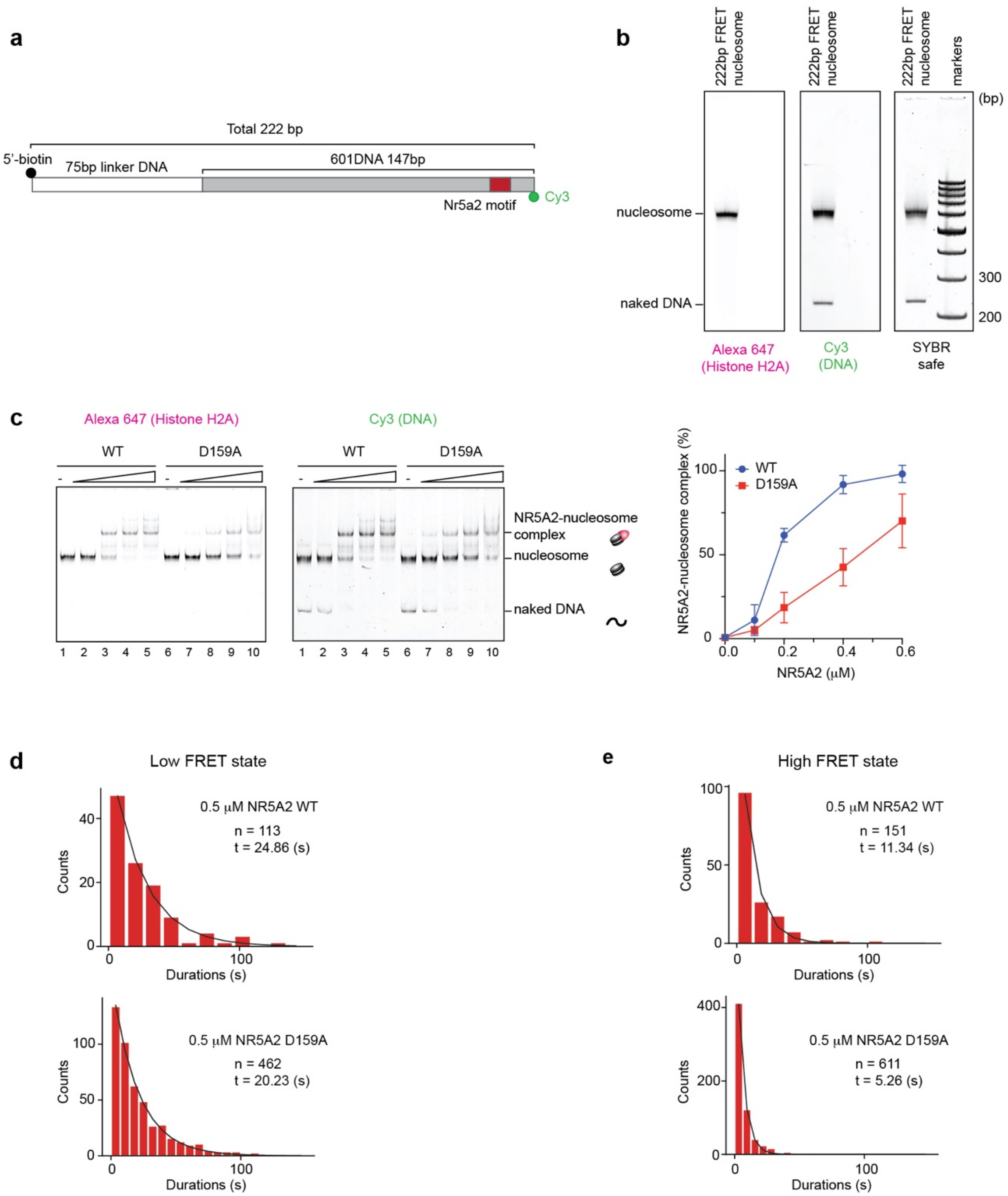


**Extended Fig. 6 DNA design for single-molecule FRET analysis**

(**a**) A 222bp of DNA used for single-molecule FRET analysis. Biotin and Cy3 fluorescence dye were conjugated each 5'-DNA end. (**b**) Purified the nucleosomes for single-molecule FRET analysis. Nucleosomes were analyzed by 6% native-PAGE and detected by SYBR gold staining and Alexa 647 fluorescence. 100 bp markers were loaded in the right side of the sample. (**c**) The left panels show representative data of EMSA with the nucleosomes used for single-molecule FRET analysis. Alexa647 fluorescence signals correspond to the nucleosome. The lanes 1-5 and lanes 6-10 show results with NR5A2 wildtype and NR5A2 D159A mutant, respectively. Right panels show the graphical representations. The average values of three independent experiments are shown with the SD values. (**d**) Histograms showing the duration in the low FRET state with the number of counts. (**e**) Histograms showing the duration in the high FRET state with the number of counts.
